# Real-World Progression-Free Survival with Erlotinib versus Osimertinib in EGFR L858R+T790M Compound Mutation Non-Small Cell Lung Cancer: An Exploratory Analysis of the MSK-CHORD Dataset

**DOI:** 10.64898/2026.06.25.734551

**Authors:** Zeinab Dalloul, Ahmad Abboud, Iman Dalloul, Mahmoud Abdelsalam

## Abstract

**Background:** Osimertinib is the standard first-line treatment for EGFR-mutant non-small cell lung cancer (NSCLC) harboring common activating mutations, including exon 19 deletions and L858R. It is also active against tumors with acquired T790M resistance. However, the EGFR L858R+T790M compound mutation — where both variants co-occur within the same tumor — may confer distinct drug-sensitivity profiles not predicted by either mutation alone. Limited data exist on comparative treatment outcomes in this rare genotype.

**Methods:** Using the MSK-CHORD clinicogenomic dataset (n=24,950), we identified patients with concurrent EGFR L858R and T790M mutations receiving erlotinib (Erlo) or osimertinib (Osi) monotherapy. Real-world progression-free survival (rwPFS) per treatment line was calculated using a strict definition requiring confirmed radiological progression events (rwPFS-strict), excluding lines with null endpoint data. Kaplan-Meier analysis, log-rank testing, Cox proportional hazards regression, and cross-cohort heterogeneity testing (Cochran’s Q statistic) were performed.

Two control cohorts — L858R-only (n=372) and T790M-only (n=76) — were analyzed in parallel to assess mutation-context specificity of treatment response.

**Results:** Thirty-one patients with EGFR L858R+T790M were identified; 21 contributed evaluable monotherapy lines, yielding 23 Erlo and 15 Osi treatment lines (14 unique patients per treatment group, 7 contributing to both). Median rwPFS numerically favored Erlo over Osi (7.10 vs 5.32 months; HR 1.29, 95% CI 0.66–2.52; log-rank p=0.46). This directional trend was reversed in the L858R-only control cohort, where Osi demonstrated significant superiority (9.03 vs 5.75 months; HR 0.70, 95% CI 0.55–0.89; p=0.003). The T790M-only cohort showed no significant difference (HR 1.32, p=0.12). An exploratory post-hoc heterogeneity test confirmed a significant cross-cohort interaction (Q=9.94, df=2, p=0.007).

**Conclusions:** The expected osimertinib advantage was absent in L858R+T790M compound-mutant NSCLC. The opposing hazard ratio directions across mutation contexts (HR 1.29 vs 0.70), with a significant exploratory cross-cohort interaction (p=0.007), suggest that the EGFR L858R+T790M compound mutation may represent a pharmacologically distinct entity with differential TKI sensitivity. These hypothesis-generating findings warrant prospective validation.

**HIGHLIGHTS:**

- L858R+T790M compound mutation may represent a distinct pharmacological TKI context.
- Erlotinib showed a point-estimate PFS advantage over osimertinib (HR 1.29, p=0.46).
- Osimertinib was significantly superior in L858R-only disease (HR 0.70, p=0.003).
- Cross-cohort HR reversal in T790M-containing cohorts; interaction p=0.007.
- Prospective validation with larger cohorts and allelic phasing data is warranted.

**TWEETABLE ABSTRACT:** #EGFR L858R+T790M compound mutation: erlotinib trend over osimertinib (HR 1.29 vs 0.70 in L858R-only), interaction p=0.007. #LungCancer

## 1. INTRODUCTION

EGFR-mutant non-small cell lung cancer (NSCLC) is driven predominantly by exon 19 deletions and the exon 21 L858R point mutation. Third-generation EGFR tyrosine kinase inhibitors (TKIs), particularly osimertinib (Osi), have supplanted first-generation agents such as erlotinib (Erlo) as the standard first-line treatment, following the FLAURA trial demonstrating superior progression-free survival,^1^ with subsequent confirmation of overall survival benefit.^2^

The T790M gatekeeper mutation in EGFR exon 20 arises in approximately 50-60% of patients as a mechanism of acquired resistance to first- and second-generation EGFR TKIs.^3,4^ Osi was developed specifically to overcome T790M-mediated resistance, with preclinical evidence demonstrating potent activity against T790M-positive cell lines,^5^ and its second-line superiority was established in the AURA3 trial.^6^ The T790M mutation increases EGFR kinase affinity for ATP, reducing first-generation TKI binding efficiency.^7^

A distinct subset of patients harbor EGFR L858R and T790M mutations as a compound mutation, with both variants detectable within the same tumor at sequencing — a rare occurrence identified in only 0.12% (31/24,950) of patients in MSK-CHORD. The clinical behavior of this rare compound mutation is poorly characterized. Regardless of whether T790M in this context is pre-existing at initial diagnosis, early clonal, or treatment-emergent, the co-occurrence of both variants may create a pharmacologically distinct TKI-binding environment. Limited published evidence, derived from studies of uncommon EGFR mutations (a category distinct from L858R, which is among the most common EGFR mutations) with co-occurring T790M, suggests that osimertinib sensitivity may be impaired when T790M is present alongside other mutations,^8^ and co-occurring T790M with common activating mutations has been associated with poor prognosis.^9^ A single case report has further described erlotinib clinical benefit with osimertinib resistance in a patient with confirmed EGFR L858R+T790M.^10^ The underlying mechanism may relate to differential drug-target binding kinetics when both mutations are simultaneously present, potentially conferring partial structural restoration of first-generation TKI binding.^11^

Here, we leveraged the MSK-CHORD real-world clinicogenomic dataset to perform an exploratory comparison of rwPFS with Erlo versus Osi in patients with EGFR L858R+T790M compound mutations, with parallel analyses in L858R-only and T790M-only control cohorts to assess mutation-context specificity.

## 2. METHODS

### 2.1. Study Design

This is a retrospective, exploratory cohort study comparing real-world progression-free survival (rwPFS) with erlotinib versus osimertinib monotherapy in patients harboring EGFR L858R+T790M compound mutations, using publicly available clinicogenomic data.

### 2.2 Data Source

The MSK-CHORD dataset (msk_chord_2024) is a real-world clinicogenomic cohort of 24,950 patients with solid tumors sequenced at Memorial Sloan Kettering Cancer Center, publicly available through cBioPortal.^12^ Clinical data including treatment history, disease characteristics, and outcomes are linked to tumor genomic profiles from next-generation sequencing. Data were accessed in April 2026.

### 2.3 Patient Population

Patients with co-occurring EGFR L858R and T790M mutations who received Erlo or Osi monotherapy were identified as the primary cohort. Erlo included its brand name Tarceva; Osi included Tagrisso. Only targeted therapy lines were analyzed; chemotherapy and immunotherapy records were excluded. Two control cohorts were constructed by computationally excluding L858R+T790M patients using patient-level identifier matching: L858R-only (T790M-negative, n=372, derived from 403 total L858R-positive patients after exclusion of the 31 compound-mutation patients) and T790M-only (L858R-negative, n=76, derived from 107 T790M-positive patients).

### 2.4 Variables and Definitions

Individual treatment records within 7 days of each other were grouped into a single treatment line; line start was defined as the earliest record start date. Combination regimens were excluded; only monotherapy lines containing exclusively Erlo or exclusively Osi were analyzed. In the primary cohort, 74.6% of target-drug lines (44/59) were pure monotherapy; the remaining 25.4% contained a target drug combined with other agents and were excluded. Disease control benefit (DCB) per treatment line was defined as stable disease, partial response, or complete response based on available clinical documentation, as captured by the DCB flag field in the MSK-CHORD dataset.

### 2.5 Endpoint Definition

Real-world progression-free survival was calculated per treatment line using a strict methodology (rwPFS-strict): progression was defined as the earliest confirmed radiological progression event or death, whichever occurred first, occurring after treatment line start. Treatment lines with progression documented on the same day as line start (rwPFS=0 days) were excluded from survival analysis. Patients without confirmed progression or death were censored at the date of last clinical assessment (defined as the last recorded clinical encounter or imaging date in MSK-CHORD). Of the 44 pure monotherapy lines identified (23 Erlo, 21 Osi), 6 were further excluded due to null rwPFS-strict data (no recorded progression or censoring date). All 6 excluded lines were osimertinib lines, yielding 38 lines (23 Erlo, 15 Osi) eligible for survival analysis. This asymmetric exclusion (0 Erlo, 6 Osi) could introduce differential selection bias if the missing Osi lines represent non-random missingness; this possibility cannot be excluded and is acknowledged as a limitation. No lines had rwPFS=0 (simultaneous progression and line start), so no additional exclusions were applied on that basis. This methodology mirrors the TreatmentScoringEngine implementation in the MSK-CHORD cBioPortal dashboard.

### 2.6 Statistical Analysis

Kaplan-Meier survival curves were estimated for each drug group. Between-group differences were assessed by log-rank test. Hazard ratios (HR) with 95% confidence intervals were estimated by Cox proportional hazards regression with Erlo as the reference (HR>1 indicates higher hazard with Osi relative to Erlo). Cross-cohort treatment-effect heterogeneity was formally assessed using Cochran’s Q statistic, computed from the log hazard ratios and their standard errors across the three cohorts; Q follows a chi-square distribution under the null hypothesis of homogeneous treatment effects (df = number of cohorts − 1 = 2). All analyses were performed in Python 3.10 using the lifelines library (v0.27) and SciPy (v1.10).^13^ Statistical significance was defined as p<0.05. Given the exploratory nature of this analysis, all p-values are reported descriptively without adjustment for multiple comparisons. Readers should note that at least seven statistical tests were performed across this analysis (three KM comparisons, Cox regression, multivariable Cox regression, DCB analysis, and the heterogeneity test), which increases the probability of false-positive findings; all results should be interpreted as hypothesis-generating.

### 2.7 Ethical Approval

This study used de-identified, publicly available data from the MSK-CHORD dataset accessible through cBioPortal. No institutional review board approval or informed consent was required.

## 3. RESULTS

### 3.1 Baseline Characteristics

Thirty-one patients with EGFR L858R+T790M were identified in MSK-CHORD. Median age at diagnosis was 69 years (IQR 63-74). The majority were female (74%), never-smokers (19/29 with available data, 66%), and presented with Stage IV disease (68%). Common metastatic sites included bone (84%), brain (61%), and liver (55%). Twenty patients received both Erlo and Osi at some point during their clinical course; 4 received Erlo only and 7 received Osi only. Additionally, 8 patients had treatment episodes with concurrent Erlo+Osi combination therapy; these lines were excluded from the monotherapy-based survival analysis per protocol (Section 2.4). Median overall survival was 25 months (IQR 13-39). A notable baseline imbalance was the TP53 co-mutation prevalence of 100% (4/4) in Erlo-only versus 43% (3/7) in Osi-only patients (Fisher p=0.10), which may influence treatment outcomes and confound group comparisons. Baseline characteristics are presented in Table 1.

**Table 1.** Baseline Characteristics of EGFR L858R+T790M Compound-Mutant Cohort (n=31)

| Characteristic | Total (n=31) | Erlo only (n=4) | Osi only (n=7) | Both (n=20) |
| --- | --- | --- | --- | --- |
| Age, median (IQR) | 69 (63-74) | 69 (68-73) | 67 (60-70) | 70 (64-75) |
| Female sex | 23 (74%) | 2 (50%) | 6 (86%) | 15 (75%) |
| Never smoker* | 19 (66%) | 2 (67%) | 5 (71%) | 12 (63%) |
| Former/current smoker* | 10 (34%) | 1 (33%) | 2 (29%) | 7 (37%) |
| White | 19 (61%) | 4 (100%) | 4 (57%) | 11 (55%) |
| Asian | 10 (32%) | 0 (0%) | 3 (43%) | 7 (35%) |
| Stage IV | 21 (68%) | 4 (100%) | 3 (43%) | 14 (70%) |
| Brain metastases | 19 (61%) | 3 (75%) | 3 (43%) | 13 (65%) |
| Bone metastases | 26 (84%) | 4 (100%) | 6 (86%) | 16 (80%) |
| Liver metastases | 17 (55%) | 3 (75%) | 4 (57%) | 10 (50%) |
| TP53 co-mutation | 20 (65%) | 4 (100%) | 3 (43%) | 13 (65%) |
| Median OS, months (IQR) | 25 (13-39) | 9 (6-16) | 14 (10-24) | 37 (19-45) |
| Deceased at data cutoff | 24 (77%) | 3 (75%) | 3 (43%) | 18 (90%) |

\*Among patients with known smoking status (n=29 total; 2 unknown). IQR, interquartile range; OS, overall survival. Erlo only = received erlotinib but never osimertinib; Osi only = received osimertinib but never erlotinib; Both = received both drugs. Of 20 patients in Both group, 19 received erlotinib before osimertinib. TP53 co-mutation prevalence: Erlo-only 100% vs Osi-only 43% (Fisher p=0.10, non-significant).

### 3.2 Primary Cohort: L858R+T790M

Of the 31 patients, 21 contributed at least one evaluable monotherapy line to the survival analysis (10 had no evaluable Erlo or Osi monotherapy lines, either because target drugs were received only as part of combination regimens or because PFS data were missing for monotherapy lines). Twenty-three Erlo monotherapy treatment lines and 15 Osi monotherapy treatment lines were eligible for analysis, contributed by 14 unique patients in the Erlo group and 14 in the Osi group, with 7 patients contributing lines to both groups; yielding 21 total unique patients in the analysis. Of the 23 Erlo lines, 22 experienced confirmed progression events and 1 was censored; all 15 Osi lines reached a confirmed progression event (0 censored). Median rwPFS was 7.10 months (95% CI 3.55-9.20) for Erlo versus 5.32 months (95% CI 1.51-7.23) for Osi. Cox regression yielded HR 1.29 (95% CI 0.66-2.52; p=0.46), indicating a non-significant directional trend toward longer rwPFS with Erlo. Kaplan-Meier curves are presented in Figure 1A. Individual patient treatment timelines are shown in Figure 2, illustrating the predominant Erlo-first sequencing pattern, concurrent Erlo+Osi combination use, and the distribution of treatment durations across 29 evaluable patients.

**Figure 1.**
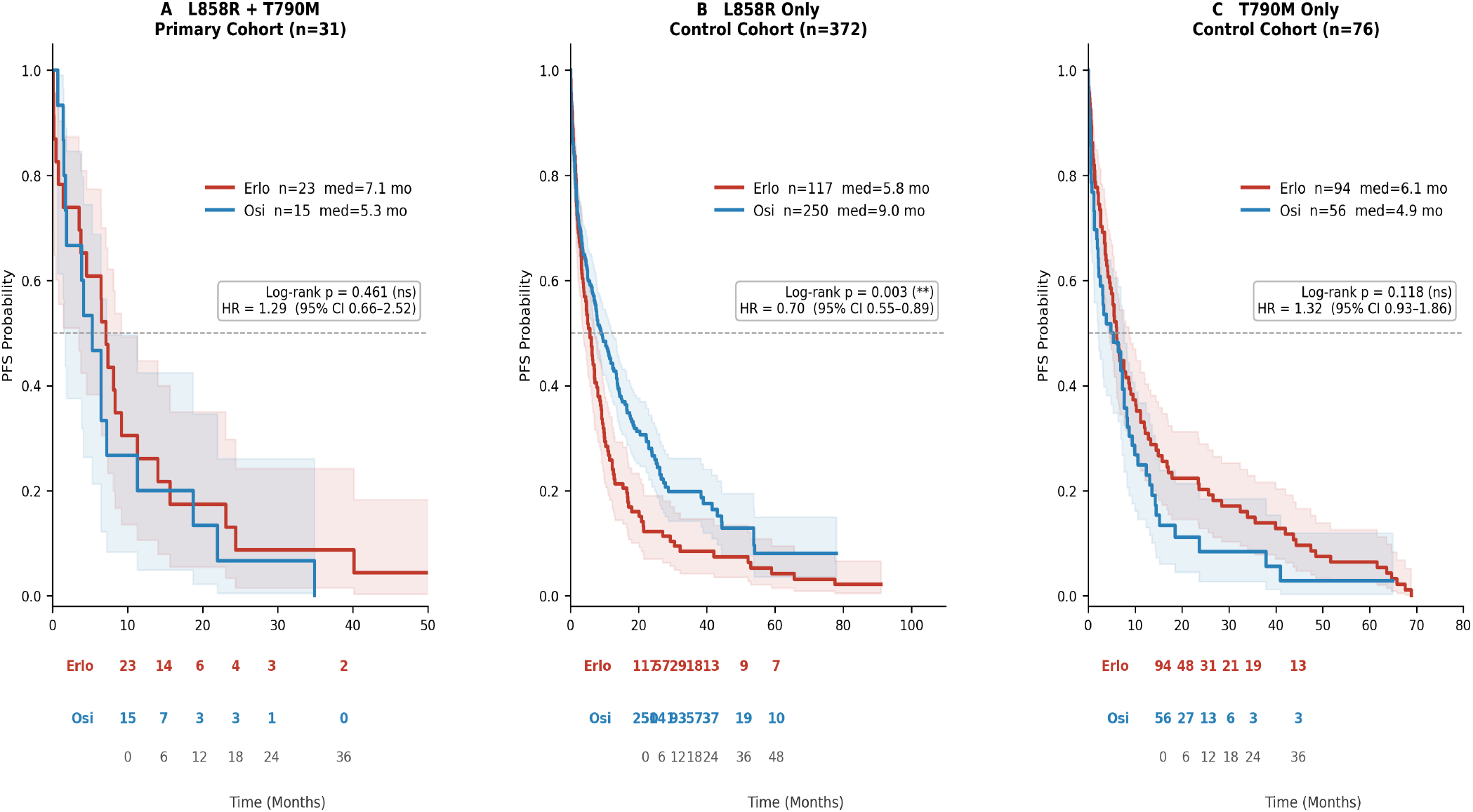
Kaplan-Meier estimates of real-world progression-free survival (rwPFS) by EGFR mutation context in the MSK-CHORD dataset. (A) Primary cohort: L858R+T790M compound mutation (n=31 patients in cohort; 23 Erlo and 15 Osi monotherapy lines from 21 contributing patients; 10 patients excluded due to combination regimens or missing PFS data). (B) Control cohort 1: L858R only, compound patients excluded (n=372 patients; 117 Erlo and 250 Osi lines). (C) Control cohort 2: T790M only, compound patients excluded (n=76 patients; 94 Erlo and 56 Osi lines). Only erlotinib (Erlo, red) and osimertinib (Osi, blue) monotherapy lines are shown. HR, hazard ratio for Osi vs Erlo from Cox proportional hazards regression (Erlo as reference); ns, not significant; *, p<0.05; **, p<0.01. Shaded regions indicate 95% confidence intervals. Number-at-risk table shown below each panel.

**Figure 2.**
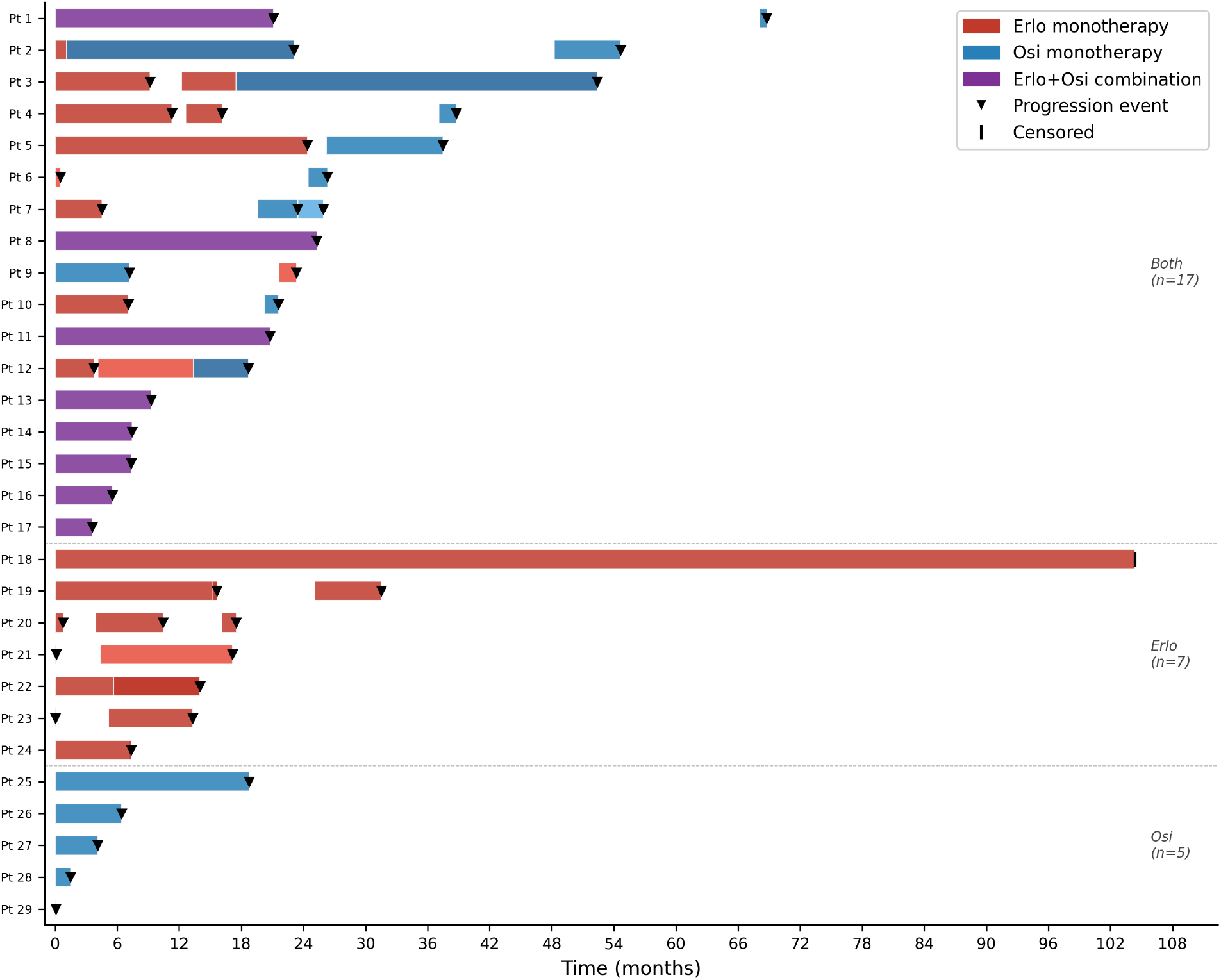
Swimmer plot of individual treatment timelines for 29 evaluable patients in the EGFR L858R+T790M cohort (2 patients excluded due to missing PFS data). Each horizontal bar represents a treatment line containing erlotinib (Erlo, red), osimertinib (Osi, blue), or concurrent Erlo+Osi combination (purple). Filled triangles denote confirmed radiological progression events; vertical bars indicate censored observations. Patients are grouped by drug exposure pattern as visible in the swimmer: both agents (n=17), Erlo only (n=7), or Osi only (n=5), and sorted by total treatment span within each group. Note: groupings differ from Table 1, which reflects overall drug history including lines not shown; 3 patients appear Erlo-only in the swimmer but received Osi outside the displayed period.

### 3.3 Control Cohort 1: L858R Only

Among 372 patients with L858R without T790M, 117 Erlo and 250 Osi monotherapy lines were analyzed. Median rwPFS was 5.75 months (95% CI 3.75-7.20) for Erlo versus 9.03 months (95% CI 6.97-12.35) for Osi (HR 0.70, 95% CI 0.55-0.89; p=0.003), with Osi demonstrating statistically significant superiority (Figure 1B). This finding is consistent with published evidence supporting third-generation TKI superiority in L858R-positive NSCLC.^1,14^

### 3.4 Control Cohort 2: T790M Only

Among 76 patients with T790M without L858R co-mutation, 94 Erlo and 56 Osi monotherapy lines were analyzed. Median rwPFS was 6.08 months (95% CI 4.57-8.80) for Erlo versus 4.86 months (95% CI 2.20-7.66) for Osi (HR 1.32, 95% CI 0.93-1.86; p=0.12), a non-significant directional trend favoring Erlo (Figure 1C).

### 3.5 Mutation-Context Specificity and Interaction Test

The hazard ratio for Osi versus Erlo differed markedly across mutation contexts: HR 1.29 (95% CI 0.66–2.52) in L858R+T790M, HR 0.70 (95% CI 0.55–0.89) in L858R-only, and HR 1.32 (95% CI 0.93–1.86) in T790M-only. In a post-hoc exploratory heterogeneity test, treatment effects were found to differ significantly across cohorts (Q=9.94, df=2, p*interaction*=0.007). This test was not pre-specified and should be interpreted as hypothesis-generating. The contrast is driven by the L858R-only cohort, where Osi demonstrated a significant advantage, in contrast to both T790M-containing cohorts (compound and T790M-only), where the hazard ratio directionally favored Erlo. This pattern is consistent with a hypothesis that co-occurring T790M may modulate osimertinib sensitivity; however, neither T790M-containing cohort independently achieves statistical significance (compound cohort p=0.46; T790M-only p=0.12), and the pattern should be considered exploratory and hypothesis-generating only.

### 3.6 Multivariable Analysis and Disease Control

In multivariable Cox regression adjusting for age, sex, smoking status, brain metastases, and TP53 co-mutation, the non-significant directional trend toward worse rwPFS with osimertinib was preserved (adjusted HR 1.53, 95% CI 0.76–3.09; p=0.24). The adjusted HR (1.53) exceeding the unadjusted estimate (1.29) suggests that measured covariates partially suppress the observed erlotinib-favoring direction, indicating that unmeasured confounding or selection bias — such as patients with poorer prognosis preferentially receiving osimertinib — may partially explain the observed effect. Female sex was the only covariate with a nominally significant association in this exploratory model (HR 0.43, 95% CI 0.19–0.98; p=0.045); however, the upper confidence bound (0.98) is very close to 1.0 and this finding should be interpreted with caution in the context of an underpowered analysis. Given the small sample size (38 treatment lines, 37 events, 6 covariates, approximately 6 events per covariate), this multivariable model is exploratory and severely underpowered; results should be interpreted as preliminary only. In a pre-specified sensitivity analysis, excluding one patient with an exceptionally long censored erlotinib line (104 months) yielded an attenuated HR of 1.14 (95% CI 0.58–2.22; p=0.72), preserving the directional finding but underscoring its fragility with respect to individual influential observations. Disease control benefit (DCB) rate per treatment line was numerically higher with erlotinib (61%, 14 of 23 Erlo lines) compared with osimertinib (47%, 7 of 15 Osi lines; Fisher p=0.51); with p=0.51, there is no significant difference in DCB rates between treatments.

## 4. DISCUSSION

In this exploratory real-world analysis, Erlo demonstrated a non-significant directional trend toward longer rwPFS compared with Osi in patients with EGFR L858R+T790M compound mutations, contrasting with the statistically significant Osi advantage in L858R-only disease. While the primary comparison did not achieve statistical significance (p=0.46) — attributable to the small sample size inherent to this rare genotype — the consistent directional HR reversal across mutation contexts (1.29 vs 0.70) is noteworthy. A post-hoc exploratory heterogeneity test confirmed significant cross-cohort interaction (p=0.007), supporting hypothesis-generating discussion of this pattern.

The EGFR L858R+T790M compound mutation — defined by co-occurrence of both variants within the same tumor at sequencing — may represent a distinct pharmacological context. A pooled analysis of osimertinib outcomes in uncommon EGFR mutations found that co-occurring T790M was associated with markedly reduced activity, with a median PFS of only 3.6 months in patients harboring multiple uncommon EGFR mutations with concurrent T790M.^8^ This pooled analysis addressed uncommon EGFR mutations — a distinct category from L858R, which is a common mutation — so direct applicability to L858R+T790M compound mutation specifically is extrapolative and should be interpreted with caution. More directly, a single case report describes erlotinib clinical benefit in a patient with confirmed EGFR L858R+T790M who demonstrated resistance to osimertinib, though anecdotal evidence from one patient cannot support broad clinical claims.^10^ Poor prognosis with T790M co-occurring with L858R has also been documented, with odds ratios of 3.37 (overall T790M+) and 3.43 (L858R+T790M subgroup specifically) for reduced recurrence-free survival.^9^ Our finding of 5.32 months with Osi in the compound-mutation cohort, compared with 9.03 months in L858R-only disease, is directionally consistent with these reports. A recent network meta-analysis found no statistically significant PFS difference between Erlo and Osi when pooled across EGFR mutation subtypes — despite Osi demonstrating superior overall survival — supporting the concept that PFS efficacy differences are mutation-subgroup dependent.^15^

Several limitations must be acknowledged. This is a retrospective observational analysis; treatment was not randomized. The primary cohort is small (n=31), limiting statistical power. Of 20 patients who received both drugs, 19 received Erlo before Osi, introducing potential sequence-driven confounding. A notable data quality concern is that all 6 monotherapy lines excluded for missing rwPFS data were osimertinib lines (0 Erlo, 6 Osi); if this missingness is non-random — for example, if Osi lines with no follow-up data represent cases of rapid progression or early discontinuation — their exclusion could artificially improve or distort Osi’s apparent survival estimate. Additionally, 8 patients received concurrent Erlo+Osi combination therapy; these lines were excluded from the primary analysis. Allelic phasing data are unavailable in MSK-CHORD; we cannot confirm whether L858R and T790M reside in cis or trans configuration, which may differentially influence drug sensitivity.^11^ Biopsy collection dates indicate that only 5 of 31 patients had genomic sequencing prior to first TKI exposure; T790M co-occurrence in the remaining 26 patients (84%) may reflect treatment-emergent acquisition rather than a pre-existing compound mutation. These two scenarios — pre-existing compound mutation at baseline versus T790M acquired under TKI pressure — represent fundamentally distinct biological contexts that may have different implications for treatment sensitivity. Future analyses should stratify by pre-treatment versus post-treatment sequencing status to distinguish these populations. However, the clinical co-occurrence of both variants at the time of sequencing is the observable and actionable feature regardless of mechanism, and we lacked the data to make this distinction reliably. As the unit of analysis was the treatment line rather than the patient, 7 patients contributed lines to both the Erlo and Osi groups; this within-patient correlation was not formally accounted for in the Cox model and may result in slightly underestimated standard errors. The number of statistical tests performed across this analysis increases the probability of false-positive findings. Finally, a numerically large but statistically non-significant imbalance in TP53 co-mutation prevalence between Erlo-only and Osi-only patient groups was noted (100% vs 43%; Fisher p=0.10); TP53 status has been associated with variable TKI outcomes and should be formally assessed in larger prospective analyses.

Despite these limitations, to our knowledge this analysis represents the largest real-world comparison of Erlo and Osi in EGFR L858R+T790M compound-mutant NSCLC to date. Prospective studies are needed to validate these findings. We recommend that such studies include: (1) pre-treatment genomic sequencing to distinguish pre-existing from treatment-emergent T790M co-occurrence; (2) allelic phasing analysis to determine cis versus trans configuration of the compound mutation; (3) formal frailty models or generalized estimating equations to account for within-patient clustering when treatment lines are used as the unit of analysis; and (4) in vitro drug sensitivity assays or patient-derived tumor models to characterize the pharmacological consequences of simultaneous L858R and T790M co-occurrence.

## 5. CONCLUSIONS

In this exploratory analysis of the MSK-CHORD dataset, erlotinib demonstrated a point-estimate median rwPFS advantage over osimertinib in patients with EGFR L858R+T790M compound mutations (7.10 vs 5.32 months), though the difference did not reach statistical significance (HR 1.29, 95% CI 0.66–2.52; p=0.46) in this underpowered analysis. This contrasts with the significant osimertinib advantage observed in L858R-only disease (p=0.003). An exploratory post-hoc heterogeneity test identified significant cross-cohort treatment-effect heterogeneity (p=0.007), and consistent HR reversal across both T790M-containing cohorts suggests that co-occurring T790M may define a pharmacologically distinct treatment context. These findings are hypothesis-generating only; prospective validation is warranted before any clinical recommendations can be made. Given the rarity of EGFR L858R+T790M (0.12% of 24,950 patients in MSK-CHORD), such validation will require multi-institutional collaboration and consortium-based genomic screening to achieve adequate sample sizes.

## DECLARATIONS

### Funding

No funding was received for this study.

### Conflicts of Interest

The authors declare no conflicts of interest.

### Author Contributions

ZD: conceptualization, methodology, writing — review and editing. AA: conceptualization, data curation, formal analysis, methodology, writing — original draft, writing — review and editing. ID: writing — review and editing. MA: clinical interpretation, writing — review and editing.

## Acknowledgments

The authors acknowledge the MSK-CHORD dataset (msk_chord_2024), publicly available via cBioPortal.

## Notes

### Competing Interest Statement

The authors have declared no competing interest.

## REFERENCES

1. Soria J-C, Ohe Y, Vansteenkiste J, et al. Osimertinib in untreated EGFR-mutated advanced non-small-cell lung cancer. N Engl J Med. 2018;378(2):113–125.

2. Ramalingam SS, Vansteenkiste J, Planchard D, et al. Overall survival with osimertinib in untreated, EGFR-mutated advanced NSCLC. N Engl J Med. 2020;382(1):41–50.

3. Kobayashi S, Boggon TJ, Dayaram T, et al. EGFR mutation and resistance of non-small-cell lung cancer to gefitinib. N Engl J Med. 2005;352(8):786–792.

4. Sequist LV, Waltman BA, Dias-Santagata D, et al. Genotypic and histological evolution of lung cancers acquiring resistance to EGFR inhibitors. Sci Transl Med. 2011;3(75):75ra26.

5. Cross DAE, Ashton SE, Ghiorghiu S, et al. AZD9291, an irreversible EGFR TKI, overcomes T790M-mediated resistance to EGFR inhibitors in lung cancer. Cancer Discov. 2014;4(9):1046–1061.

6. Mok TS, Wu Y-L, Ahn M-J, et al. Osimertinib or platinum-pemetrexed in EGFR T790M-positive lung cancer. N Engl J Med. 2017;376(7):629–640.

7. Yun CH, Mengwasser KE, Toms AV, et al. The T790M mutation in EGFR kinase causes drug resistance by increasing the affinity for ATP. Proc Natl Acad Sci USA. 2008;105(6):2070–2075.

8. Hu S, Wang C, Wang C, Zhao K, Wang Z, Dong W. Insensitivity to T790M mutation? A pooled analysis of outcomes following osimertinib for the treatment of NSCLC patients harboring uncommon EGFR mutations. Front Pharmacol. 2022;13:986962.

9. Gao X, Zhao Y, Bao Y, Yin W, Liu L, Liu R, et al. Poor prognosis with coexistence of EGFR T790M mutation and common EGFR-activating mutation in non-small cell lung cancer. Cancer Manag Res. 2019;11:9621–9630.

10. Zhao X, Dong H, Wang Z, et al. EGFR primary T790M and L858R double mutation confers clinical benefit to erlotinib and resistance to osimertinib in one lung adenocarcinoma patient. J Cancer Sci Ther. 2018;10(11):366–370.

11. Tian P, Wang Y, Wang W, Li Y, Wang K, Cheng X, et al. High-throughput sequencing reveals distinct genetic features and clinical implications of NSCLC with de novo and acquired EGFR T790M mutation. Lung Cancer. 2018;124:205–210.

12. Jee J, Fong C, Pichotta K, et al. Automated real-world data integration improves cancer outcome prediction. Nature. 2024;636:728–736.

13. Davidson-Pilon C. lifelines: survival analysis in Python. J Open Source Softw. 2019;4(40):1317.

14. Takeyasu Y, Yoshida T, Masuda K, Matsumoto Y, Shinno Y, Okuma Y, et al. Distinct progression and efficacy of first-line osimertinib treatment according to mutation subtypes in metastatic NSCLC harboring EGFR mutations. JTO Clin Res Rep. 2024;5(2):100636.

15. Runzer-Colmenares FM, Ruiz R, Maco L, Maldonado M, Puma-Villanueva L, Galvez-Nino M, et al. Comparison of erlotinib vs. osimertinib for advanced or metastatic EGFR mutation-positive NSCLC without prior treatment: a network meta-analysis. Cancers. 2025;17:1895.

